# Neuromechanical Simulation with NEURON and MuJoCo

**DOI:** 10.1101/2025.06.17.660217

**Authors:** Chris Fietkiewicz, Linh Tran, Robert McDougal, Clayton Jackson, Roger D. Quinn, Hillel J. Chiel, Peter J. Thomas

## Abstract

In computational neuroscience, simulation platforms generally do not have adequate tools to model the brain, body and environment simultaneously. We demonstrate a method for simulating neuromechanical models using a novel combination of widely used software platforms: NEURON and MuJoCo. Different neural models are used to control a realistic musculoskeletal model in both open-loop and closed-loop configurations. Three models are presented: (1) an open-loop model using simple spiking neurons from the NEURON model library; (2) an open-loop model using realistic, spiking motoneurons; and (3) a closed-loop central pattern generator with feedback from the physics engine.

## 1 INTRODUCTION

Modern computational neuroscience has a wide variety of sophisticated simulation tools for modeling the nervous system. However, neural simulation platforms generally do not have adequate tools to simulate the physical world, which is crucial since the brain is embodied and the body is situated in a complex environment [7]. Some previous studies have addressed this need by integrating general physics engines with well known neural simulation platforms. For example, Gazebo is a robotics simulator [22] that has been used by different research groups in combination with the NEST neural simulator [2, 5, 11, 12, 32]. In some cases, platforms have been developed to facilitate the integration of Gazebo with NEST, such as MUSIC [9], the Neurorobotics Platform [26], and EBRAINS [10]. In contrast to Gazebo, some modelers have used the OpenSim simulator [28], which provides specialized features for biomechanics, such as musculotendon dynamics [2, 18, 20, 25]. Many other publicly available physics simulators exist, especially for robotics applications (see [8] for a review). The MuJoCo simulator [30] is widely used in the fields of robotics and machine learning [1, 8]. Studies have reported the performance of MuJoCo to be as much as 600 or 900 times faster than OpenSim [17, 31].

Here we extend previous work using the NEURON platform to incorporate biomechanics [13]. NEURON is a widely used neural simulation platform that has enjoyed continuous development and active support for over three decades [4, 6, 14, 15, 23, 24]. NEURON is readily extensible and has a large and active user community. We demonstrate a novel combination of NEURON with MuJoCo using Python. As a preliminary effort, we used relatively small neural models with a realistic musculoskeletal model in both open-loop and closed-loop configurations. The goal for all models was to produce oscillatory motion in a musculoskeletal model of a rat hind limb. Three models were developed: (1) an open-loop model using simple spiking neurons from the NEURON model library, (2) an open-loop model using realistic, spiking motoneurons, and (3) a closed-loop central pattern generator with feedback on muscle length from the physics engine.

## 2 METHODS

The overall simulation technique consisted of a Python control program with a loop that synchronously advanced the separate simulators, NEURON and MuJoCo, by a single numerical integration timestep (dt = 0.025 ms). Before each integration step, muscle forces computed from the NEURON simulator were used to set force parameters for actuators in the MuJoCo simulator. For closed-loop models, muscle lengths computed in MuJoCo were used to set parameters in NEURON. All communication between Python and the two simulators was done using the standard API for each platform.

All simulations used the same MuJoCo musculoskeletal model of a rat hind limb. The model is based on skeletal components from a publicly available OpenSim model that included a hip, femur, tibia, and foot [19]. Components from the OpenSim model were converted to a format compatible with MuJoCo v2.3.5, including STL format for geometries and XML format for 3-dimensional placement of the geometries. MuJoCo flexor-extensor tendons were added and positioned following analysis of moment arms. Each MuJoCo tendon was controlled by an actuator with a parameter for applied force that was set programmatically using the Python control program. A single joint was located at the top of the femur, while the tibia and foot remained in fixed positions relative to the femur. There were no surface contacts, and the leg hung vertically from the hip while at rest.

For all models, neural and muscle components were implemented using NEURON v8.2. All models included two neurons, each innervating a single muscle that controlled either leg protraction or retraction, respectively. Two open-loop models were studied with different neuron types in order to demonstrate the versatility of the NEURON simulator. They used a physiological muscle model from Fietkiewicz et al. [13], which had been adapted from Kim and Heckman [21]. The first open-loop model used a Hodgkin-Huxley neuron [16] from the standard NEURON model library. The second open-loop model used a more realistic motoneuron, based on Purvis and Butera [27], that can exhibit a much lower spike rate as compared to a Hodgkin-Huxley neuron. Figure 1(a) shows the overall design for both open-loop models.

**Fig. 1.**
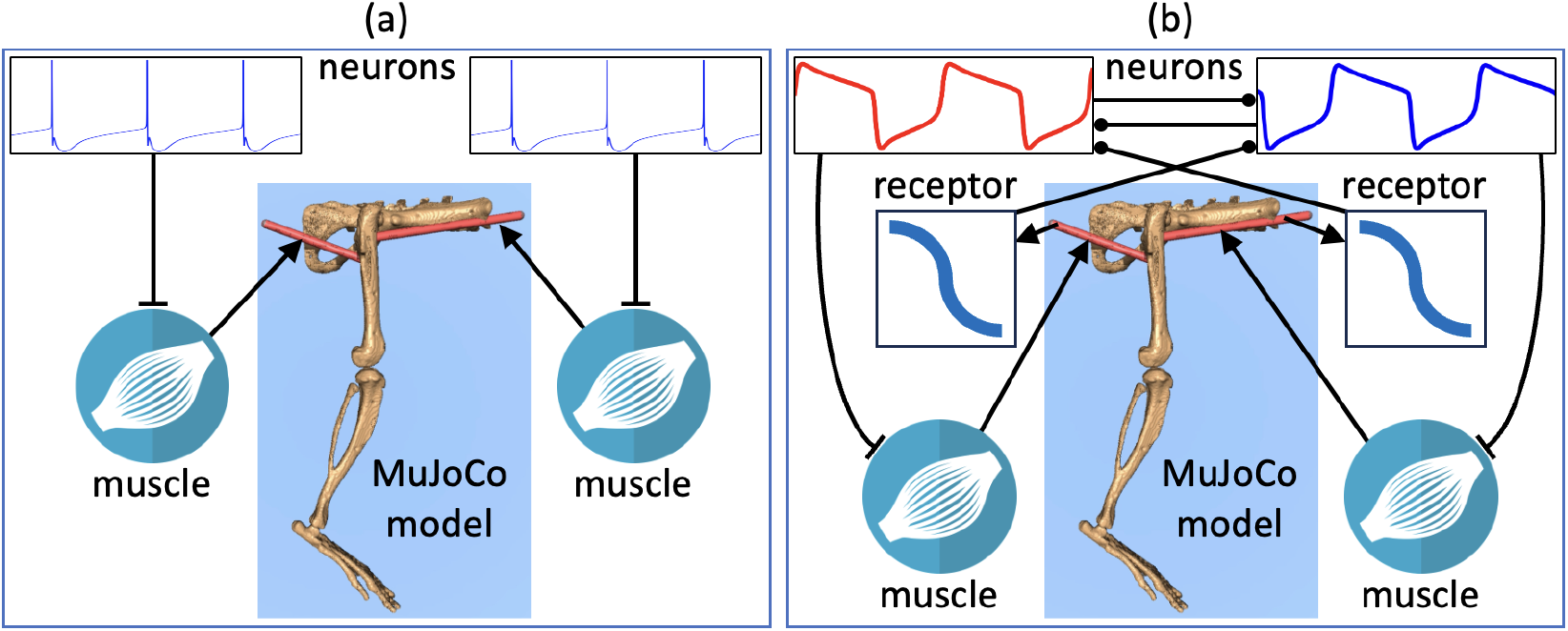
Model designs. (a) Open-loop model. (b) Closed-loop model. Neurons and muscles were implemented in NEURON. Synaptic connections included inhibitory (circles) and excitatory (perpendicular lines). The MuJoCo model included a skeleton and tendons (shown in red).

The closed-loop model was based on a model by Yu and Thomas [33] and later adapted for NEURON [13]. The key features included two conductance-based neurons that form a central pattern generator (CPG) and receive sensory feedback from muscle stretch receptors. Each neuron receives inhibitory input from the other neuron and the sensory receptor for the contralateral muscle. Previous versions of the model [13, 33] computed muscle length based on the reciprocating movement of a 1-dimensional pendulum. In the present work, muscle forces were applied to actuators in the MuJoCo model to move the leg. A key feature is that muscle length, calculated in MuJoCo, is a parameter that is used to compute the stretch receptor feedback used in the NEURON model. Figure 1(b) shows the overall design for the closed-loop model.

## 3 RESULTS

For both open-loop models, each of the two neurons received a stimulus pulse at a different start time than the other, with periods that did not overlap, in order to produce a leg retraction followed by protraction. The stimuli differed for each of two open-loop models due to a difference in spike rates and the resulting rate of increase in force produced by the muscle. For the first model, containing Hodgkin-Huxley neurons [16], stimuli had a 10 nA amplitude, a 70 ms pulse duration, and a 200 ms offset between pulse start times. For the second model, containing motoneurons [27], stimuli had a 1 nA amplitude, a 360 ms pulse duration, and a 400 ms offset between pulse start times.

Figure 2 shows the behavior of the neural, muscular, and skeletal components for both models, and a rendered animation is available [3]. It can be seen that the membrane voltage of each neuron exhibits action potential “spikes” during stimulation, resulting in a gradual increase in corresponding muscle force. After neuron #1 stops spiking, calcium dynamics maintain the applied force before the force returns to zero Newtons. During this phase, the angle of the skeleton leg increases proportionally from rest in synchronization with the force of muscle #1 for retraction. Neuron #2 begins spiking shortly before the force of muscle #1 has decreased to zero. During this period of overlap in applied forces, an increasing force in muscle #2 counteracts the retraction force, and the leg angle returns to zero when the two opposing muscle forces are equal. The leg then begins a protraction phase where the negative leg angle is proportional to the force of muscle #2. The differences in time scale between the responses of the two models is due to the different neuron spike rates.

**Fig. 2.**
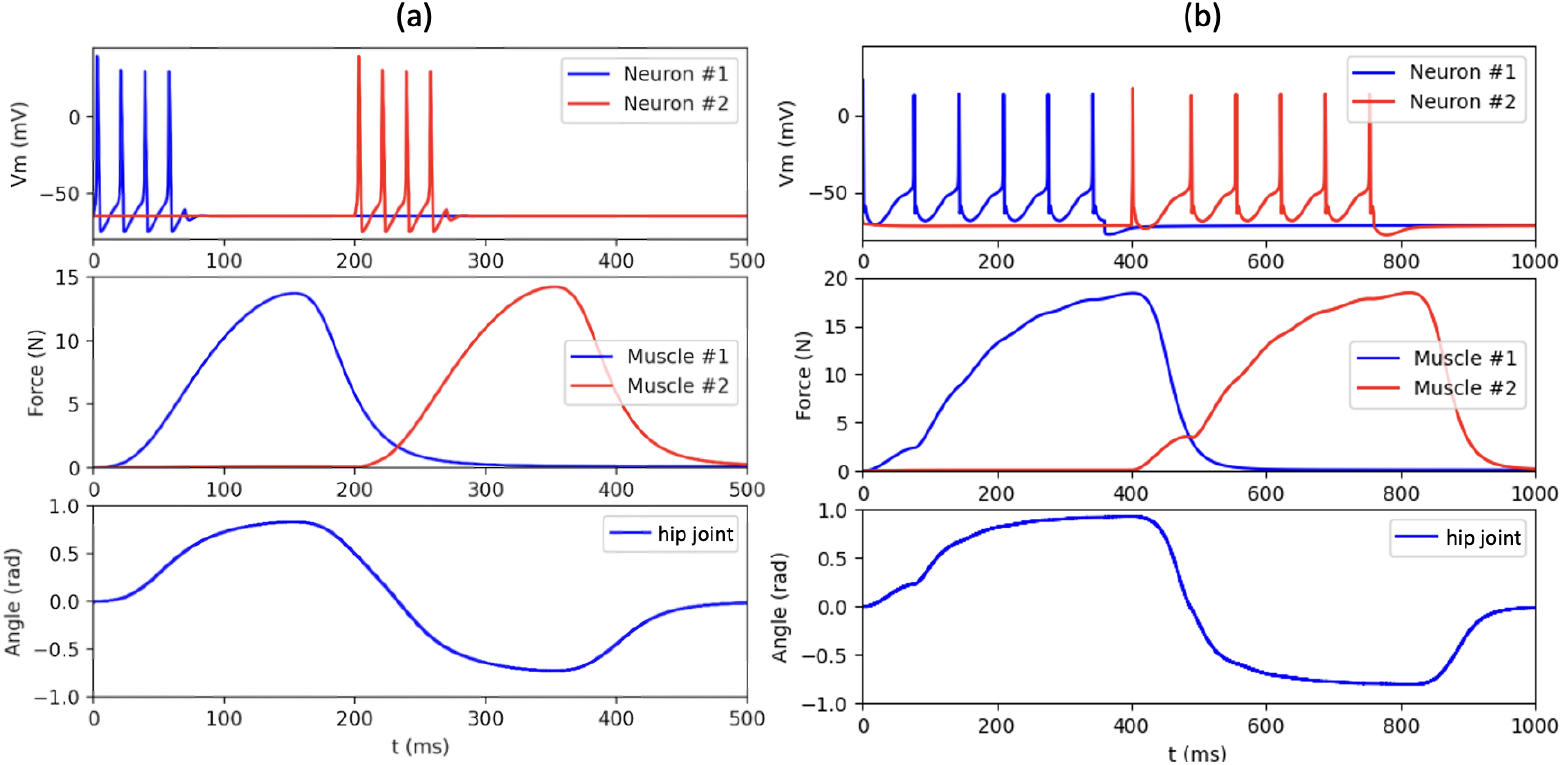
Simulation of two open-loop models: (a) Model which uses Hodgkin-Huxley neurons. (b) Model which uses motoneurons. Each simulation shows neuron voltages (top), applied force (middle), and hip joint angle (bottom). An animation is available online [3].

In contrast to the previous open-loop models, the closed-loop model maintains an oscillatory rhythm without external stimulation because of internal dynamics, as has been previously studied using muscle and mechanical models that were different than those used here [33]. The results of the present model are very similar to those in [13, 33]. Figure 3(a) shows two complete cycles of activity, with a single cycle lasting approximately 3,570 ms, and a rendered animation is available [3]. Note that the time axes in Figure 3(a) begin at t = 640 ms in order to omit initial transitory behavior due to the bidirectional interconnectivity between neural, muscular, and skeletal components.

**Fig. 3.**
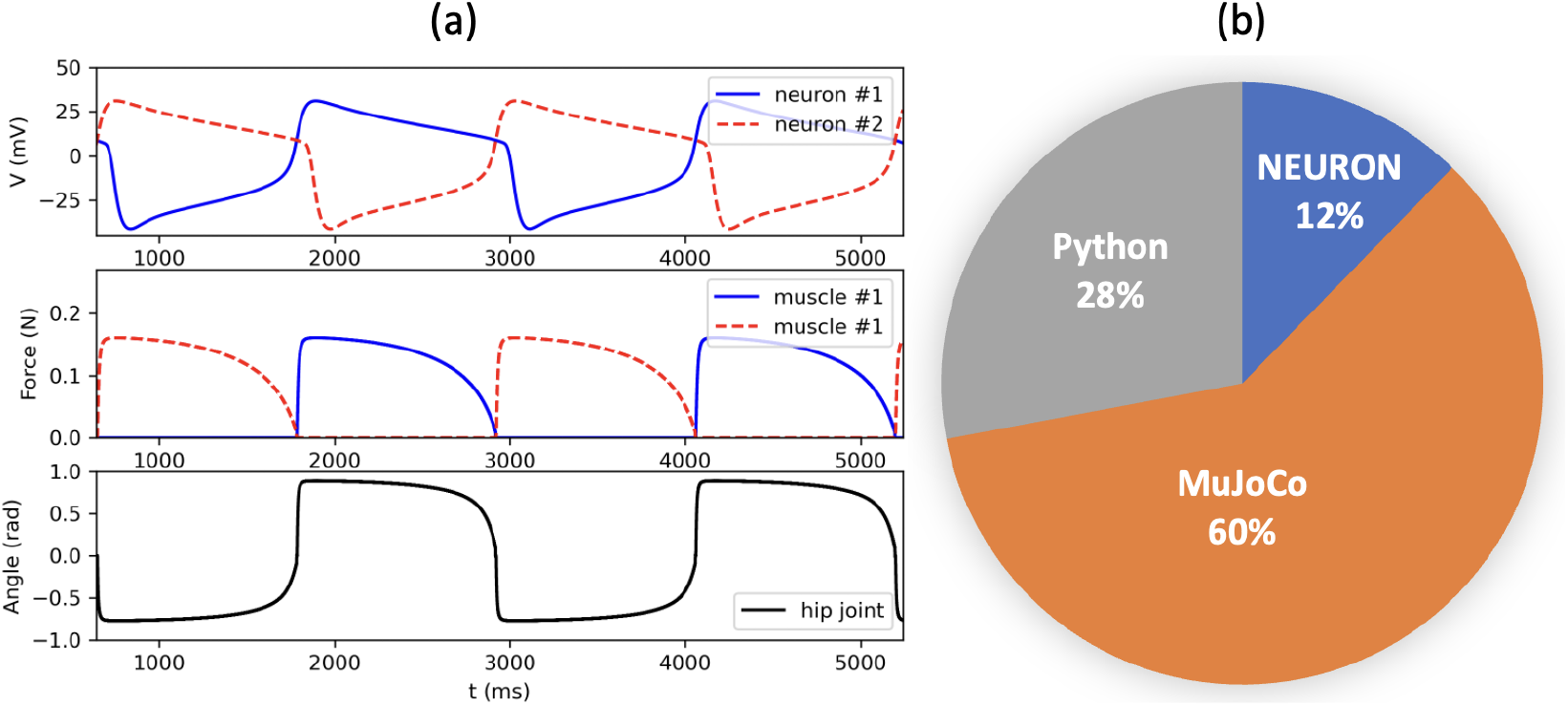
Closed-loop model. (a) Simulation results show neuron voltages (top), applied force (middle), and hip joint angle (bottom). (b) Relative computation time for software platforms used during simulation. An animation of the model is available online [3].

We analyzed the computation time of the three software platforms used for the simulation, including the use of Python for the main control loop which reads and writes parameter values in the two simulator platforms. The analysis was done on a MacBook Air with an M2 processor and MacOS 14.0. For a simulation time of 10,000 ms, the total computation time for the simulation loop was an average of 12.7 sec. The relative time for each software platform was determined by selectively disabling the MuJoCo numerical integration and parameter reading and writing. The analysis is shown in Figure 3(b), where the results, in order of greatest to least, were the MuJoCo simulator (60%), Python control (28%), and the NEURON simulator (12%).

## 4 CONCLUSION

To address the need for simulating neuromechanical models, we demonstrated a novel combination of the widely used software platforms NEURON and MuJoCo. Different neural models were used to control a realistic musculoskeletal model in both open-loop and closed-loop configurations. The models used several different features available in NEURON, as well as a relatively simple Python control program to exchange parameters with MuJoCo. In an analysis of computational time, MuJoCo required the largest amount, followed by the Python control program. Future work should investigate methods for increasing efficiency. For the Python control program, alternatives may include more efficient techniques for memory access and the use of a Python compiler. Additionally, MuJoCo supports a C++ API that may offer an alternative control strategy. Ultimately, this technique will facilitate quantitative comparisons of motor control models for organisms with differences in limb length and locomotion cycle period [29].

## AUTHOR CONTRIBUTIONS

CF, HJC, and PJT conceived the research. CF and LT carried out the research. HJC and PJT supervised the research. CF, RM, HJC, and PJT wrote the paper. CJ and RDQ developed the MuJoCo model.

## ACKNOWLEDGMENTS

We thank Zhuojun Yu for assistance in modifying the CPG model. This work was supported in part by National Institutes of Health BRAIN Initiative grant RF1 NS118606-01, National Institutes of Health grant R01 NS011613-46, National Science Foundation grant DBI 2015317 as part of the NSF/CIHR/DFG/FRQ/UKRI-MRC Next Generation Networks for Neuroscience Program, and the Oberlin College Department of Mathematics.

## Notes

### Competing Interest Statement

The authors have declared no competing interest.

